# Processing *of Clostridium perfringens* Enterotoxin by Intestinal Proteases

**DOI:** 10.1101/2025.02.11.637699

**Authors:** Archana Shrestha, Jessica L. Gonzales, Juliann Beingesser, Francisco A. Uzal, Bruce A. McClane

**Author notes:** Address Correspondence to: Bruce A. McClane, Room 427 Bridgeside Point II Building, 450 Technology Drive, Pittsburgh, PA 15219. **Key Contribution:** During diseases, *C. perfringens* enterotoxin (CPE) is released into the intestinal lumen, which is rich in proteases. The study determined that, i) CPE processing involving trypsin occurs rapidly *in vitro*, *ex vivo* and in the intestines, ii) this processed CPE is present in CPE large pore complexes and iii) this CPE retains cytotoxic and enterotoxic activity.

## Abstract

*C. perfringens* type F isolates are a leading cause of food poisoning and antibiotic-associated diarrhea. Type F isolate virulence requires production of *C. perfringens* enterotoxin [CPE], which acts by forming large pore complexes in host cell plasma membranes. During disease, CPE is produced in the intestines when type F strains undergo sporulation. The toxin is then released into the intestinal lumen when the mother cell lyses at the completion of sporulation. Once present in the lumen, CPE encounters intestinal proteases. This study examined *in vitro*, *ex vivo* and *in vivo* processing of CPE by intestinal proteases and the effects of this processing on CPE activity. Results using purified trypsin or mouse intestinal contents detected rapid cleavage of CPE to a major band of ∼32 kDa and studies with Caco-2 cells showed this processed CPE still forms large complexes and retains cytotoxic activity. When mouse small intestinal loops were challenged with CPE, the toxin caused intestinal histologic damage despite rapid proteolytic processing of most CPE to 32 kDa within 15 min. Intestinal large CPE complexes became more stable with longer treatment times. These results indicate that CPE processing involving trypsin occurs in the intestines and the processed toxin retains enterotoxicity.

## Introduction

*Clostridium perfringens* is a major human gastrointestinal pathogen [1–6]. Annually, this bacterium is estimated to cause one or five million food poisoning cases in, respectively, the USA or European Union [3, 7]. The vast majority of *C. perfringens* human foodborne disease cases are caused by type F strains [8], with type F food poisoning ranking as the second most common bacterial foodborne disease in the USA [3, 7]. Type F strains also cause 5-15% of all cases of nonfoodborne human gastrointestinal [GI] disease, most notably including antibiotic-associated diarrhea [9–11].

Production of *C. perfringens* enterotoxin [CPE], a single polypeptide of 35 kDa, is essential for type F strain enteric virulence [12]. In human enterocyte-like Caco-2 cells, CPE action starts when this toxin binds to receptors, which include certain members of the claudin family of ∼20-27 kDa proteins located in epithelial or endothelial tight junctions [13–15]. Binding of CPE to those claudin receptors forms an ∼90 kDa small complex that contains CPE plus both a receptor and a nonreceptor claudin [16]. Approximately 6 CPE small complexes oligomerize into an ∼425-500 kDa large complex that initially corresponds to a prepore [16, 17]. Each CPE molecule sequestered in this prepore then extends a β-hairpin to create a β-barrel pore that is selectively permeable to cations, including calcium[18–20]. With prolonged treatment times, a secondary large CPE complex of ∼550-660 kDa containing CPE, receptor and nonreceptor claudins, and the tight junction protein occludin can sometimes be detected in Caco-2 cells [21]. Calcium influx into CPE pore-containing Caco-2 cells then activates calpain [20], which kills those cells via either classical caspase-3 mediated apoptosis [when a lower CPE dose treatment causes limited calpain activation] or necroptosis [when a higher CPE dose treatment causes strong calpain activation] [22, 23].

In the small intestine, CPE also binds to claudin receptors on villus tip cells, followed by formation of a large CPE complex [24, 25]. The ability to form this large CPE complex also appears to be important for CPE-induced intestinal damage based upon results from studies using several CPE variants blocked at various steps in CPE action [24, 26]. Those experiments revealed that only CPE variants capable of forming large complexes in Caco-2 cells were able to produce intestinal histologic damage in mouse small intestinal loops. Other studies have strongly suggested this CPE-induced intestinal histologic damage is responsible for intestinal fluid and electrolyte losses that clinically manifest as diarrhea [27, 28].

During GI disease, CPE is produced when type F isolates sporulate in the intestines [1, 9, 29]. Once made, CPE is not secreted but, instead, accumulates in the cytoplasm of the mother cell [28, 29]. When sporulation is completed, the mother cell lyses to free its mature spore, which concomitantly releases CPE into the intestinal lumen [28]. After discharge, CPE encounters an environment rich in intestinal proteases, including trypsin and chymotrypsin but also other proteases.

Many bacterial protein toxins produced in the intestines, including several *C. perfringens* toxins [1], are cleaved by intestinal proteases. A number of previous studies reported that, *in vitro,* CPE can also be processed by purified trypsin or chymotrypsin [30–33]. Specifically, purified trypsin was shown to remove the first 25 amino acids [∼3 kDa] from the N-terminus of CPE to generate an ∼32 kDa CPE fragment. Similarly, purified chymotrypsin reportedly removes the first 36 N-terminal amino acids from CPE, which should generate a CPE fragment of ∼31 kDa [30].

However, the intestinal lumen contains multiple proteases [34] and the extent of CPE processing, if any, under intestinal conditions has not yet been examined despite its potential disease relevance given that proteolytic cleavage affects the toxicity of other *C. perfringens* toxins [1]. Therefore, this study evaluated the effects of mouse small intestinal content [MIC] on CPE proteolytic processing and compared those *ex vivo* results to the *in vitro* effects of purified trypsin or chymotrypsin on CPE to discern the involvement of those two proteases in CPE proteolytic processing by intestinal contents. Studies were then extended to evaluate CPE proteolytic processing and toxicity in the intestines using mouse small intestinal loops; these analyses also provided new insights regarding the formation kinetics and stability of the large CPE complex *in vivo*.

## Results

### Further examining purified trypsin or chymotrypsin effects on CPE processing in vitro

The current study first sought to confirm previous reports [31–33] that, *in vitro*, purified trypsin reduces the molecular mass of CPE by ∼3 kDa. Consistent with those reports, a slight [∼3 kDa] reduction in CPE molecular mass was detected [Fig. 1A] by either CPE western blotting or Coomassie blue staining of 12% acrylamide gels after electrophoresis of toxin that was incubated with purified trypsin for 10 min at 37°C. The extent of this rapid CPE processing did not increase even using a 10-fold higher trypsin concentration. CPE western blotting of those samples revealed that this *in vitro* trypsin treatment caused a reduction in immunoreactivity relative to CPE incubated in the absence of trypsin. Since the level of CPE present after trypsin treatment showed little decrease upon Coomassie blue staining and previous studies [31, 33] reported that trypsin removes N-terminal CPE sequences *in vitro*, these results indicate that the N-terminal CPE sequences removed by purified trypsin contribute to one or more epitopes recognized by the CPE polyclonal antiserum used in this study.

**Fig. 1.**
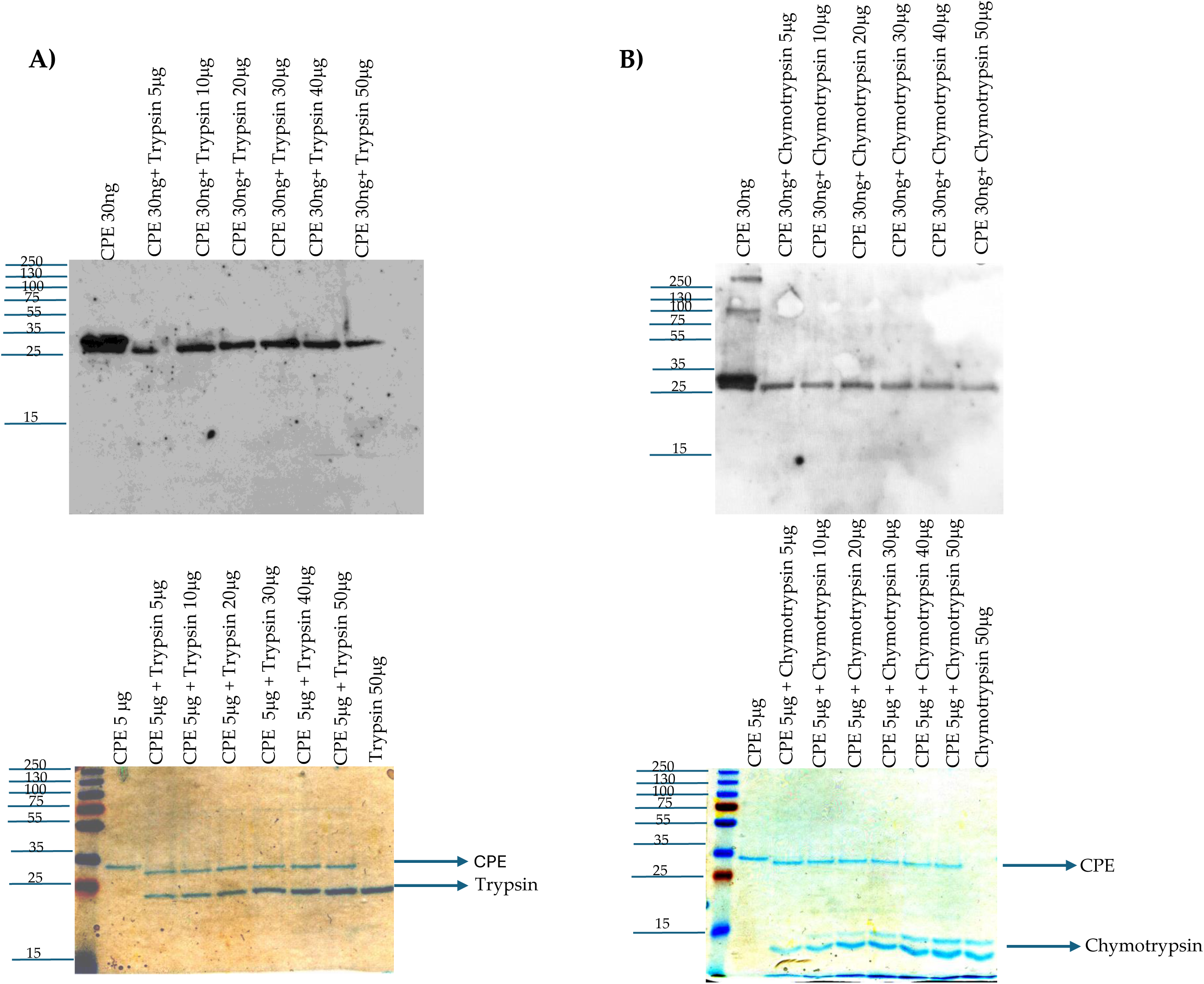
Effects of purified trypsin or chymotrypsin on CPE processing *in vitro*: A] Trypsin treatment. Top panel: 30 ng of CPE was incubated with 0, 5, 10, 20, 30, 40, or 50 μg of trypsin in PBS buffer for 10 min at 37°C. After this incubation, samples were electrophoresed without boiling on a 12% acrylamide gel containing SDS and then CPE western blotted. Bottom panel: 5 μg of CPE was treated with 0, 5, 10, 20, 30, 40, or 50 μg of trypsin in PBS buffer for 10 min at 37°C. The samples were then electrophoresed as described above, followed by staining with Coomassie blue [bottom panel]. B] Chymotrypsin treatment. Top panel: 30 ng of CPE was treated with 0, 5, 10, 20, 30, 40, or 50 μg of chymotrypsin in PBS buffer for 10 min at 37°C. After incubation, samples were electrophoresed without boiling on a 12% acrylamide gel containing SDS and then CPE western blotted. Bottom panel: 5 μg of CPE was treated with 0, 5, 10, 20, 30, 40, or 50 μg of chymotrypsin in PBS buffer for 10 min at 37°C. The samples were then electrophoresed as described above, followed by staining with Coomassie blue [bottom panel]. Numbers at left of gels show the size of protein standards by molecular mass [in kDa]. Results shown in both panels are representative of three repetitions for each gel.

A previous amino acid sequencing study [30] reported that purified chymotrypsin cleaves native CPE at multiple amino acid residues and concluded this processing mainly produces a CPE fragment with the first 36 N-terminal sequences removed. However, that previous study never evaluated the size or relative abundance of the major CPE fragment[s] generated by treatment of CPE with purified chymotrypsin *in vitro*. Therefore, the current study performed both CPE western blotting and Coomassie blue staining after SDS-PAGE of samples containing CPE incubated with or without purified chymotrypsin [Fig. 1B]. After a 10 min incubation at 37°C, Coomassie blue staining of those gels showed that purified chymotrypsin had rapidly processed CPE to a single band only slightly [∼1 kDa] smaller than CPE. The extent of this processing did not increase using even a 10-fold higher chymotrypsin concentration. CPE western blotting of similar gels found similar results and also detected a reduction in immunoreactivity for this CPE fragment vs. CPE after chymotrypsin treatment. Since the level of CPE present after chymotrypsin treatment showed little decrease when visualized by Coomassie blue staining, the CPE region removed by chymotrypsin must also contribute to one or more epitopes recognized by the CPE polyclonal antiserum used in this study.

### Effects of mouse intestinal content [MIC] on CPE processing ex vivo

Since intestinal contents contain a mix of trypsin, chymotrypsin and other proteases [34], this study next examined the effects of MIC from 3 different mice on CPE processing *ex vivo*. When analyzed by CPE western blotting, these three MIC all rapidly and similarly processed CPE to a major species that had only a small [∼3 kDa] reduction in molecular mass compared to native CPE [Fig. 2A]. As observed in Fig. 1 using purified trypsin or chymotrypsin, the intensity of the CPE western blot signal decreased upon MIC treatment.

**Fig. 2.**
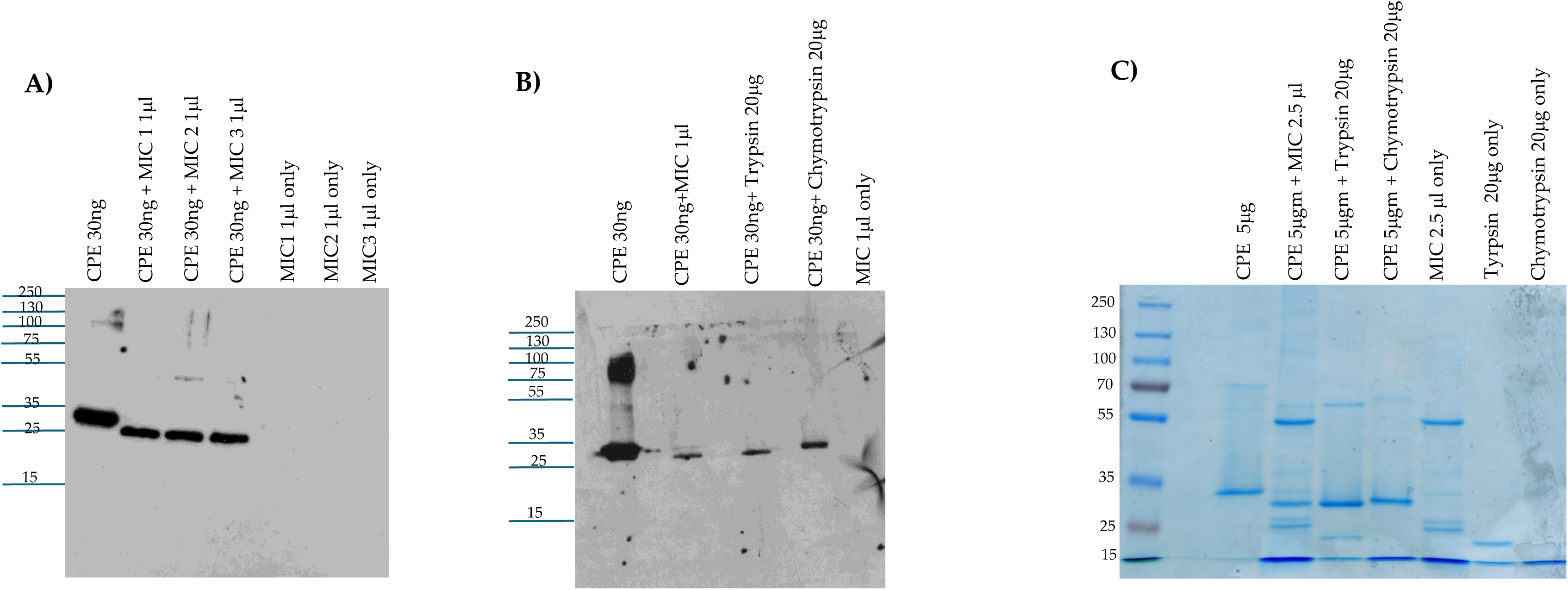
Effects of mice intestinal contents on CPE: A] MIC effects on CPE processing. MIC were prepared from 3 different mice [2 female and 1 male]. To evaluate their effects on CPE, 30 ng of native CPE was or was not mixed with 1 μl of each MIC in PBS and incubated at 37 °C for 10 mins. After this incubation, samples were boiled, electrophoresed on a 12% acrylamide gel and then processed for CPE western blot analysis. The same amount of each MIC alone [no CPE] in PBS was similarly CPE western blotted to assure that the observed immunoreactivity was due to the presence of CPE. B] Head-to-head CPE western blot comparison of the effects of MIC, purified trypsin or chymotrypsin on CPE processing. MIC alone [no CPE] was also included as a specificity control. Before electrophoresis and CPE western blotting as described for panel A, MIC or MIC plus CPE samples, but not CPE plus trypsin or chymotrypsin samples, were boiled. C] Head-to-head Coomassie blue staining comparison of the effects of MIC, purified trypsin or chymotrypsin on CPE processing. Samples were prepared and electrophoresed as in panels A and B except that 5 μg of CPE and/or 2.5 μl of MIC were added to samples [when specified]. Also electrophoresed were equivalent concentrations of MIC, purified trypsin or purified chymotrypsin in the absence of CPE, for comparison. Samples with MIC were boiled but other samples were not. Panel A, B and C results are a representative result of three repetitions. Numbers at left of gels or blots indicate size of standard proteins [in kDa].

The next CPE western blot experiment [Fig. 2B] compared the head-to-head migration of CPE on 12% acrylamide gels containing SDS after the toxin had been incubated with purified trypsin, purified chymotrypsin, or MIC. This analysis revealed that *ex vivo* incubation of CPE with MIC generated a CPE fragment that migrates similarly as the CPE fragment produced by *in vitro* incubation of CPE with purified trypsin. By comparison, the CPE fragment generated by treatment with purified chymotrypsin in Fig. 2B migrated more slowly than the trypsin- or MIC-generated CPE fragments on the same western blot. These results provide support for the involvement of trypsin in CPE processing by MIC. On those Fig. 2B western blots, there was again some decrease in CPE species signal intensity after treatment with MIC.

When this experiment was repeated with Coomassie blue staining [Fig. 2C], several protein bands were visible in the MIC alone samples [no CPE present]. Subtracting those background MIC bands from lanes run with samples where both CPE and MIC were present together supports similar CPE processing as detected by western blotting, i.e., the major CPE fragment was ∼3 kDa smaller after MIC processing. These results were also consistent with similar CPE processing by MIC and trypsin. Finally, compared to CPE alone, there was only a slight decrease in staining intensity for the MIC-generated CPE fragment in Fig. 2C; combining that result with the strong decrease in western blot signal intensity visible in Fig. 2A and 2B after MIC treatment indicates that, as with purified trypsin or chymotrypsin treatment, MIC disrupt one or more epitopes in CPE.

### Effects of trypsin- or chymotrypsin-specific inhibitors on CPE processing by MIC

To investigate further the involvement of trypsin in the MIC-generated CPE processing shown in Fig. 2, the trypsin-specific inhibitor TLCK was employed. To verify the specificity of this inhibitor, we first showed [Fig. 3A] that, i] preincubating purified trypsin with TLCK blocked CPE processing, but ii] this inhibitor did not block CPE processing by purified chymotrypsin.

**Fig. 3.**
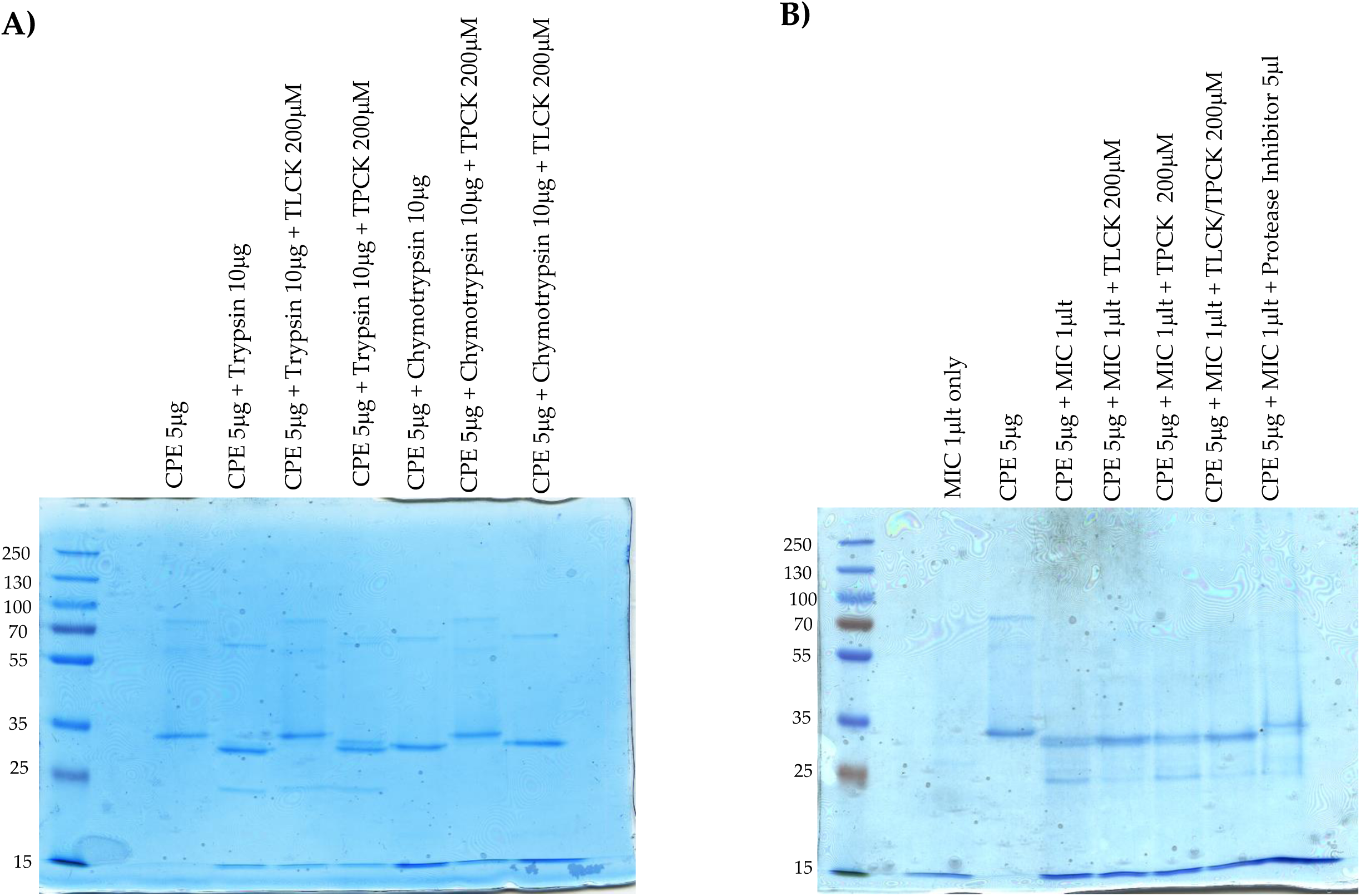
Effects of protease inhibitors on CPE processing. Panel A] 10 μg of purified trypsin was preincubated for 30 min at 37°C with or without 200 μM TLCK or 200 μM TPCK in PBS; 10 μg of purified chymotrypsin was similarly preincubated with or without 200 μM TPCK or 200 μM TLCK in PBS. After this preincubation, 5 μg of CPE was added to each sample and those samples were further incubated for 10 min at 37°C. The incubated samples were then electrophoresed on 12% acrylamide gels containing SDS; after electrophoresis the gels were stained with Coomassie blue. Panel B] 1 μl of MIC was preincubated for 30 min at 37°C with, i] 200 μM TLCK, ii] 200 μM TPCK iii] 200 μM of both TCPK and TLCK, iv] 5 μl of resuspended Protease Inhibitor Cocktail III in PBS. As a control, 1 μl of MIC was similarly preincubated in PBS only. After preincubation, 5 μg of CPE was added to each samples and the samples were further incubated for 10 min at 37°C. The incubated samples were then boiled and electrophoresed on 12% acrylamide gels containing SDS, followed by staining the gels with Coomassie blue. Results shown are representative of three repetitions.

For these Fig. 2 studies, Coomassie blue staining of gels was performed to detect any protease-generated CPE fragments that might lack epitopes and thus would not be visualized by CPE western blotting. In addition, to simplify these analyses, less MIC volume was used than for Fig. 2C experiments to avoid the presence of detectable levels of MIC proteins upon Coomassie blue staining, as confirmed by results of the MIC alone lane shown in Fig. 3B results, where no protein bands were visible. With the MIC preparations used in Fig 3B, a major CPE band ∼3 kDa less than CPE was again detected, along with a minor CPE band reduced in size by ∼7 kDa.

Preincubating MIC with TLCK prior to the addition of CPE caused a slight increase in size of the main CPE fragment and a distinct increase in staining intensity of this CPE fragment compared to the main CPE fragment generated by MIC in the abscence of TLCK [Fig 3B]. In addition, a decrease in staining intensity was noted for the MIC-generated minor CPE band when TLCK was present; this reduction in minor CPE band levels likely explains the increased staining intensity for the prominent CPE band in Fig. 3B samples when both MIC and TLCK were present. These Fig. 3B results indicated that TLCK can effectively impact normal CPE processing by MIC and thereby further support the involvement of trypsin in CPE processing by MIC.

Similar experiments were repeated using the chymotrypsin inhibitor TPCK. Preincubation of purified chymotrypsin with TPCK blocked subsequent CPE processing, while preincubation of purified trypsin with TPCK had no subsequent effect on CPE processing, verifying inhibitor specificity [Fig. 3A]. Preincubation of this MIC preparation with TPCK still produced a major ∼32 kDa CPE fragment similar to that produced by MIC alone [no inhibitor] and a minor ∼28 kDa CPE band was also visible at the same staining intensity as with MIC alone [Fig 3B]. Therefore, these results do not provide support for a chymotrypsin requirement in normal MIC processing of CPE.

Conceivably, chymotrypsin effects on CPE processing might become detectable in the absence of trypsin activity, so MIC were also preincubated with both TPCK and TLCK inhibitors. Compared to the effects of trypsin inhibitor alone, no difference in MIC processing of CPE was observed in the presence of both inhibitors. Lastly, the observation of some CPE processing in the presence of both TLCK and TPCK suggested that other MIC proteases can also process CPE in absence of trypsin. Consistent with that possibility, pan-protease inhibitors completely blocked CPE processing by MIC [Fig. 3B].

### Analysis of MIC effects on large CPE complex formation in Caco-2 cells

As described in the Introduction, evidence indicates that large CPE complex formation is important for CPE pore formation, CPE-induced cytotoxicity and CPE-induced intestinal damage. Therefore, the current study next assessed [Fig. 4A] whether *ex vivo* exposure of CPE to MIC affects subsequent formation of large CPE complexes in human enterocyte-like Caco-2 cells. For this evaluation, CPE was added to Caco-2 cells that were incubated in either HBSS containing MIC or HBSS alone; lysates of those cells were then electrophoresed on 6% acrylamide gels containing SDS to detect large CPE complex formation [large CPE complexes are SDS-resistant [35]]. In the absence of MIC, large CPE complex formation was detectable at the 15 min time-point and the amount of this complex then increased further between 15 and 30 min of that treatment. Importantly, the presence of MIC had no effect on large CPE complex formation levels at either time-point.

**Fig. 4.**
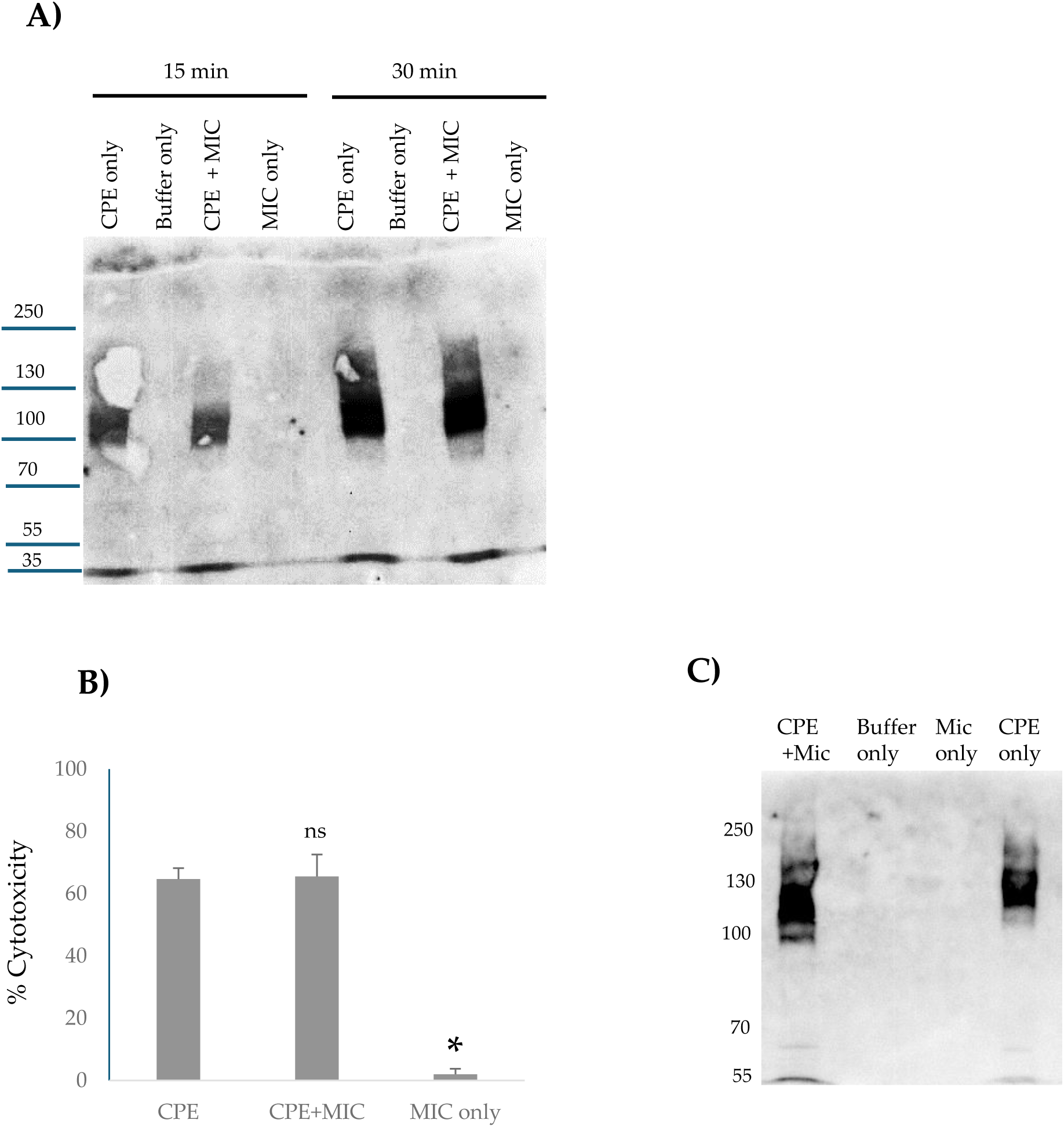
Effects of MIC on large CPE complex formation and CPE-induced cytotoxicity in Caco-2 cells: A] MIC effects on large CPE complex formation. MIC in HBSS was added to confluent Caco-2 cells, followed immediately by addition of 0.5 μg/ml of CPE for 15 or 30 min. Other control wells received the same amount of MIC in HBSS [no CPE] or HBSS alone. After incubation the cells were harvested gently and lysed with RIPA buffer in the presence of Benzonase and Protease Inhibitor cocktail at RT with gentle rotation for 30 min. Lysates were centrifuged and 1 μl of each supernatant was mixed with 24 μl of HBSS plus 5 μl 5X loading buffer. That mixture was loaded without boiling onto a 6% acrylamide gel with SDS. After electrophoresis and transfer, blots were processed for CPE Western blotting. Results shown are representative of three western blots. Numbers at left of gel indicate size of standard proteins [in kDa]. B] Effects of preincubating CPE with MIC on CPE-induced cytotoxicity. Before their addition to Caco-2 cells, HBSS containing 3 μg of CPE, 3 μg of CPE with 2.5 μl of MIC, 2.5 μl of MIC, or HBSS buffer alone were preincubated at 37°C for 10 min. The samples were then collected and adjusted with HBSS to 1 μg/ml of CPE, or diluted similarly for MIC alone, and added onto confluent Caco2 cells for 1 h at 37°C. After incubation, the samples were gently collected and spun down and the lysates were processed for LDH release, as a marker for cytotoxicity, using the CyQUANT LDH Cytotoxicity Assay Kit. Results shown are the means of 3 repetitions. Error bars show SD. The cytotoxicity difference between CPE vs. CPE plus MIC was not statistically significant [p<0.05], but * indicates that treatment with MIC only was significantly [p<0.05] different from CPE treatment. C] Effects of preincubating CPE with MIC on large CPE complex formation by panel B Caco-2 cells. Panel B Caco-2 cells were lysed and processed for CPE western blotting as described in Panel A. Results shown are representative of three repetitions.

### Effects of preincubating CPE with MIC on CPE-induced cytotoxicity in Caco-2 cells

We next evaluated [Fig. 4B] whether MIC effects on CPE processing impact CPE-induced cytotoxicity in Caco-2 cells. This experiment involved preincubating CPE with HBSS that did or did not contain MIC *ex vivo* for 10 min at 37°C [as in Fig.2] and then applying those samples to Caco-2 cells for 1 h at 37°C. Results of this experiment indicated that preincubating CPE with MIC did not significantly affect the subsequent cytotoxic activity of CPE for Caco-2 cells [Fig 4B]. As a control, a 1 h treatment of Caco-2 cells with HBSS containing the same concentration of MIC alone [no CPE] did not induce cytotoxicity, indicating that the cytotoxicity observed in Caco-2 cells treated with both MIC and CPE was attributable to the toxin. Consistent with these cytotoxicity results, CPE western blotting [Fig. 4C] detected similar levels of large CPE complex formation in those same Caco-2 cells, whether they were treated with CPE that had or had not been preincubated with MIC.

### Analyses of large CPE complex formation kinetics and stability in the mouse small intestine

We previously reported [25] that CPE also forms a large complex in mouse small intestine. However, the time-course of large CPE complex formation *in vivo*, as well as any stability changes to that intestinal large CPE complex over time, have not been examined. While lacking, information regarding the kinetics and stability of large CPE complex in the intestines would be useful for designing and interpreting experiments to assess proteolytic processing of CPE *in vivo*. Therefore, the current study first CPE-challenged mouse small intestinal loops for 15 min, 30 min or 4 h [Fig. 5]. Intestinal contents from those mice then were [to promote CPE dissociation from complex] or were not boiled, followed by CPE western blotting after electrophoresis on 6% acrylamide gels containing SDS. In these CPE western blot analyses, immunoreactive material was present in mouse small intestinal loops treated with CPE in buffer, but not in loops treated with buffer alone, confirming the specificity of the western blot for CPE detection.

**Fig. 5.**
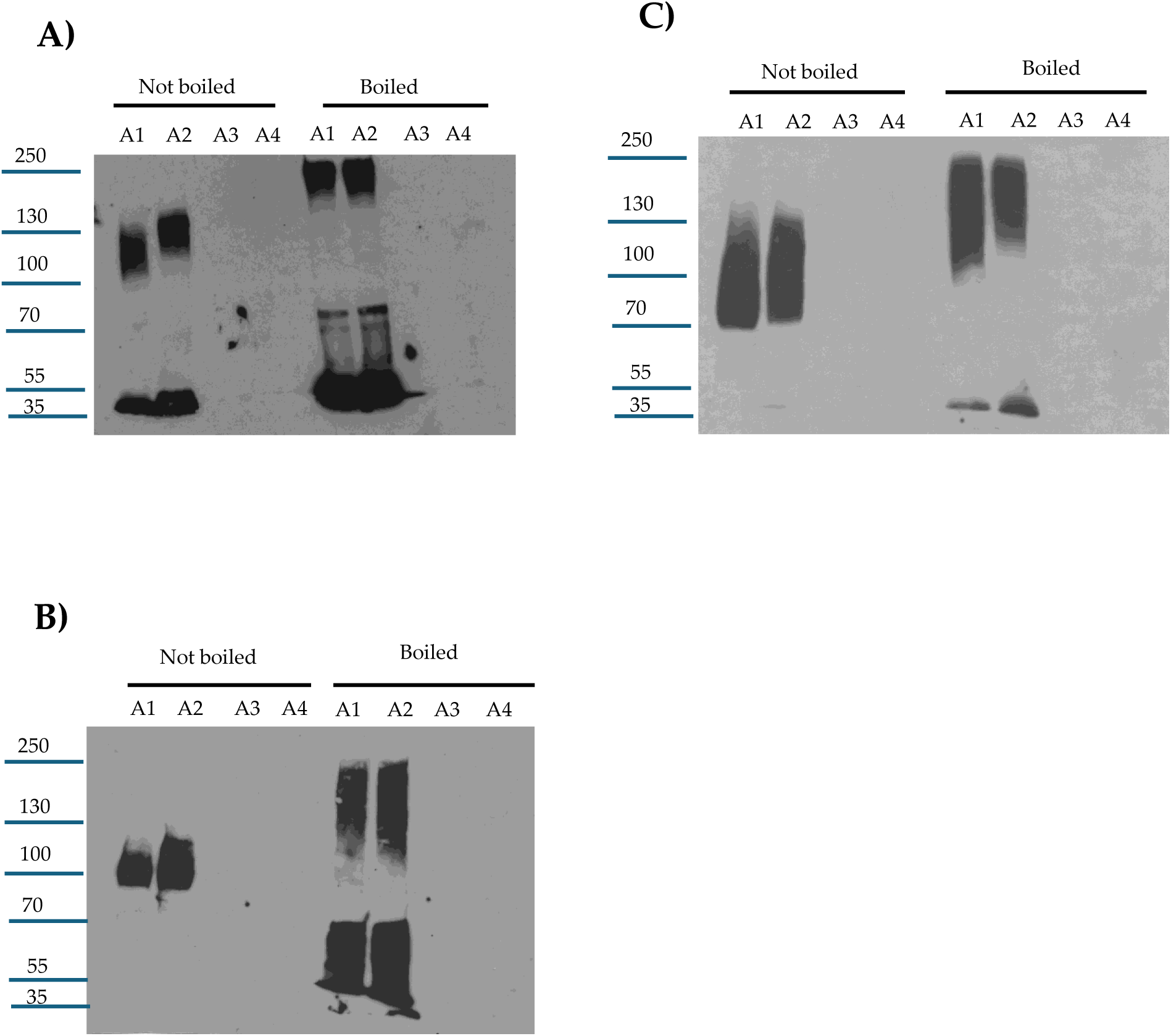
Analyses of *in vivo* large CPE complex formation kinetics and large CPE complex stability. Mouse small intestinal loops were treated with HBSS containing 100 μg/ml of CPE or HBSS alone for 15 min, 30 min or 4 h. The small intestinal contents were then collected, electrophoresed on 6% acrylamide gel with SDS, and processed for CPE western blot analysis with or without sample boiling [as indicated]. A] 15 min treatment B] 30 min treatment and C] 4 h treatment. Note that large CPE complex migrates anomalously on 6% acrylamide gels with SDS [16], so its migration vs. standard proteins [as indicated on left of gel in kDa] does not accurately reflect the size of this complex. Shown are representative western CPE blots for three repetitions for each time-point.

This experiment [Fig. 5A] detected appreciable *in vivo* formation of large CPE complex after only 15 min of CPE-treatment of mouse small intestinal loops, indicating that both CPE binding [which is required for large CPE complex formation [36]] and large CPE complex formation begin quickly in the intestines. However, even without sample boiling, substantial amounts of noncomplexed CPE were also detectable in CPE western blots of those 15 min time-point intestinal content samples. This noncomplexed CPE could be free, unbound CPE present in the lumen and/or CPE bound to the intestines in CPE small complex, which dissociates in SDS [35]. When the 15 min intestinal content samples were boiled, the ratio of free to large CPE complex-associated toxin increased, indicating some CPE had dissociated from the large CPE complex.

Similar western blot analyses were also performed with mouse small intestinal loops treated for 30 min [Fig. 5B] or 4 h [Fig. 5C] with the same CPE dose. For intestinal contents from loops treated with CPE for 30 min, the ratio of free CPE to large complex-associated CPE in nonboiled samples was much lower than for samples from loops treated with CPE for 15 min, indicating that nearly all CPE present in those 30 min CPE treatment samples was now associated with the large CPE complex and, by extension, nearly all CPE had bound to the intestines by 30 min. However, boiling those 30 min samples still caused the appearance of substantial free CPE on these western blots, i.e., similar to the samples from mice treated with CPE for 15 min, boiling induced significant dissociation of CPE from large CPE complex in intestinal contents from mice subjected to a 30 min CPE treatment.

Like the 30 min CPE treatment results, little free CPE was detected in nonboiled samples from loops treated with the toxin for 4 h. However, in contrast to the 15 min or 30 min CPE treatment results, boiling caused only limited CPE dissociation from the large CPE complex present in mouse intestinal contents from loops treated for 4 h with CPE. This indicates that the large CPE complex formed in the small intestine physically changes over time to become more stable, i.e., more resistant to dissociation by heat and SDS.

### Evaluating whether the large CPE complex formed in CPE-treated mouse small intestine contains full-length or processed CPE

Since Fig. 5 confirmed rapid large CPE complex formation in mouse small intestinal loops, it was important to distinguish whether the CPE associated with this *in vivo* large CPE complex had been proteolytically-processed by intestinal proteases or if it was still intact due to rapid intestinal binding and/or by protection from proteases once sequestered within the large complex. This question was addressed by treating mouse small intestinal loops with CPE for 15 min, 30 min or 4 h and then subjecting those samples to CPE western blotting on i] a 6% acrylamide gel with SDS [no sample boiling] to confirm large CPE complex formation in these samples or ii] on a 12% SDS-containing polyacrylamide gel with sample boiling in order to promote complex dissociation to better visualize the size of CPE fragments.

Fig. 6A results confirmed Fig. 5 results by showing that large CPE complex formation became detectable by 15 min of loop challenge. After 30 min of loop challenge, the vast majority of CPE was associated with large CPE complex. The Fig. 6B experiments then provided new information indicating that, even after only 15 min of CPE treatment, much of the toxin bound to the intestines had already been proteolytically-processed. Specifically, on CPE western blots of these 12% acrylamide gels with SDS and sample boiling, much of the CPE in the intestines electrophoresed as a smaller band than native CPE and co-migrated with CPE processed *ex vivo* by MIC. Similar to Fig. 3 *ex vivo* results, a minor fraction of the dissociated CPE had been processed further to a size ∼7 kDa less than CPE at all time-points evaluated in this experiment. Since nearly all CPE present in the 30 min or 4 h samples, and a substantial portion of the CPE in 15 min samples, was associated with large CPE complex, these results indicate that much of the CPE present in the large complex formed in mouse small intestinal loops has been proteolytically-processed.

**Fig. 6.**
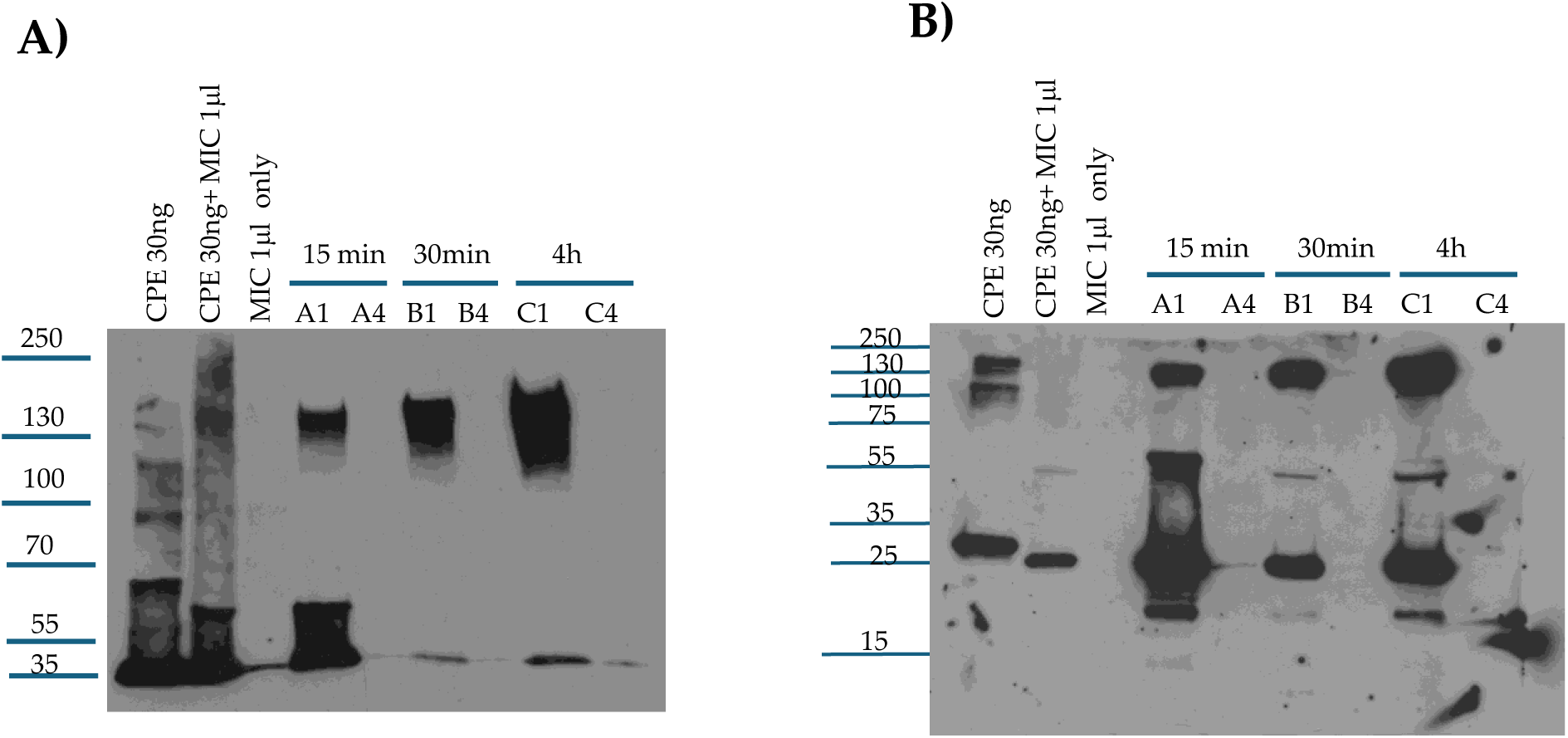
Analysis of CPE processing *in vivo*. A] Confirmation of large CPE complex formation in mouse small intestinal loops. Loops A1, B1 and C1 received 100 μg/ml of CPE in HBSS for 15 min, 30 min or 4 h respectively. Loops A4, B4 and C4 received only HBSS [no CPE] for 15 min, 30 min or 4h, respectively. Intestinal contents were then electrophoresed, without boiling, on a 6% acrylamide gel with SDS and CPE western blotted. Markers at left of blot show migration of protein standards in kDa although, as mentioned in the Fig. 5 legend, the large CPE complex migrates anomalously on 6% acrylamide gels with SDS, so its migration vs standard proteins does not accurately reflect its true size. Also shown is migration of CPE alone, CPE treated with MIC *ex vivo* and MIC alone for comparison. B] Analysis of CPE size after treatment of mouse small intestinal loops. After loops were treated as described for panel A. Intestinal contents were then electrophoresed, with sample boiling, on 12% acrylamide gels with SDS before CPE western blotting. Markers at left of blot show migration of protein standards in kDa. Also shown is migration of CPE alone, CPE treated with MIC *ex vivo* and MIC alone for comparison. Gels shown in Panels A and B are representative of three repetitions.

### Assessing whether the proteolytically-processed CPE associated with large CPE complex formed in the small intestine retains enterotoxicity

Since Fig. 6 represents the first definitive indication that much of the CPE associated with the large CPE complex formed in the intestines undergoes proteolytic-processing, histologic analyses were performed on those intestines to confirm this processed CPE retains enterotoxicity, i.e., the ability to cause intestinal damage.

Those histologic analyses detected little or no intestinal damage after a 15 min CPE challenge of mouse small intestinal loops. However, by 30 min of CPE challenge, a time by which much of the CPE in the loops had been processed and was almost completely associated with large CPE complex [Fig. 6], CPE began to cause small intestinal damage, including villus blunting and epithelial desquamation [Fig. 7]. This damage became more severe by 4 h of CPE treatment [Fig. 7].

**Fig. 7:**
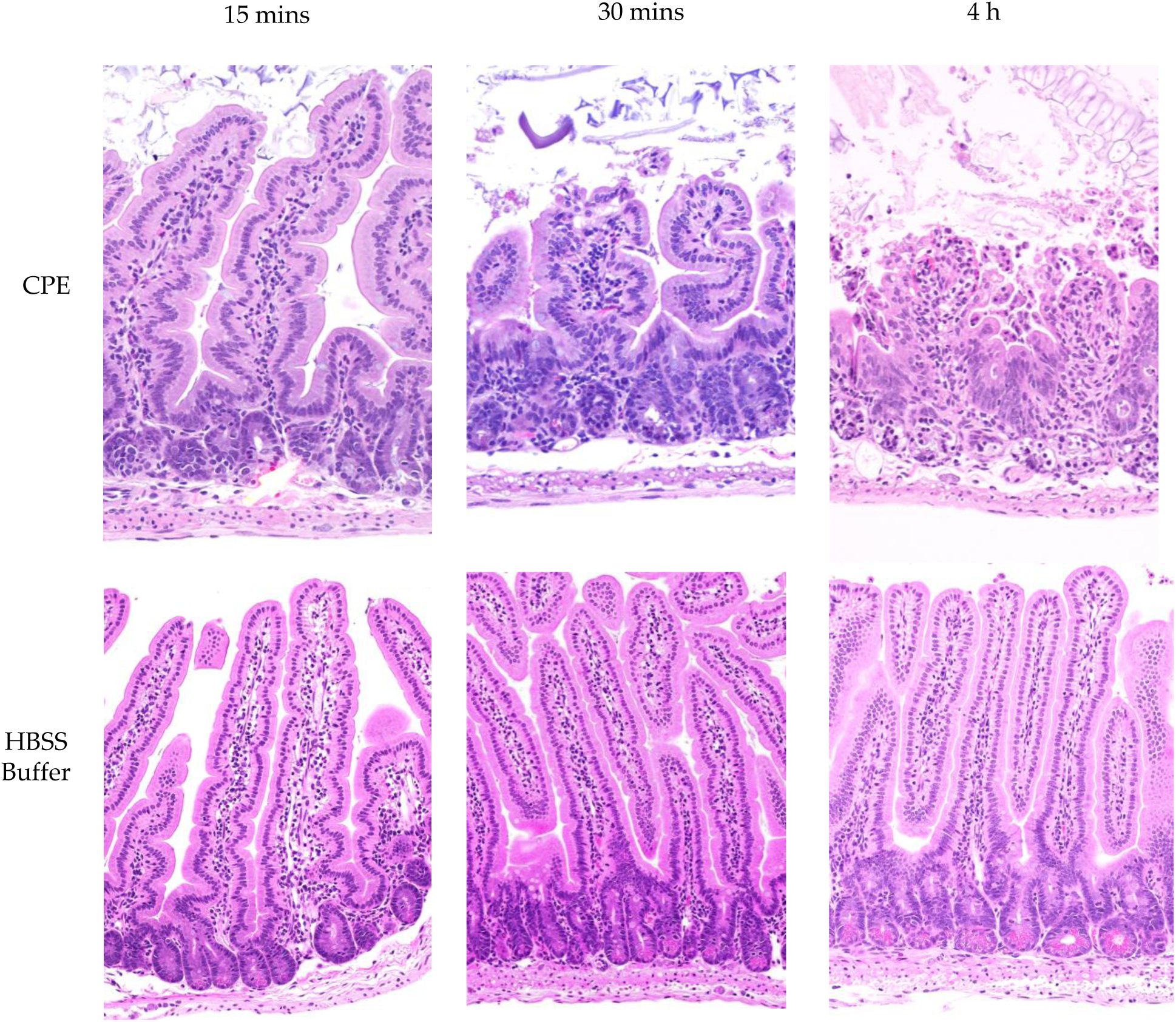
CPE processed in the small intestines retains enterotoxicity. Mouse small intestinal loops were challenged with HBSS containing 100 μg/ml of CPE [top row] or HBSS alone [bottom panel] for 15 min, 30 min or 4 h, as indicated. After these treatments, samples from the intestinal loops were processed routinely for the production of 4 μm thick hematoxylin and eosin [HCE] stained sections. Results shown are representative of 3 mice for each challenge.

## Discussion

Intestinal proteases process several *C. perfringens* toxins produced in the intestines, e.g., beta, iota and epsilon toxins [1, 37–40]. Previous studies [30–33] had examined the *in vitro* effects of purified trypsin or chymotrypsin on CPE and reported that those proteases cleave CPE. Using amino acid sequencing, mass spectrometry and western blot approaches, several studies [31–33, 41] reported that purified trypsin removes the first 25 N-terminal amino acids [∼3 kDa] from CPE. The current SDS-PAGE results using purified trypsin are consistent with this conclusion. The current study also determined that trypsin treatment decreased western blot immunoreactivity using a polyclonal CPE antiserum. This result indicates the first 25 amino acids of CPE contribute to one or more epitopes, which is consistent with a previous epitope mapping study [32] that showed a CPE_25-319_ fragment lost a non-neutralizing epitope present in native CPE.

Another amino acid sequencing study [30] concluded that purified chymotrypsin removes the first 37 N-terminal amino acids from CPE; however, this conclusion has been confusing for two reasons. First, that previous study also detected chymotrypsin cleavage at several N-terminal amino acids preceding CPE amino acid 37, although it did not evaluate the size or relative predominance of CPE fragment[s] produced by chymotrypsin cleavage *in vitro*. Second, the same group earlier reported [41] that native CPE is resistant to chymotrypsin cleavage without reduction and S-carboxylation. In response, the current study evaluated purified chymotrypsin cleavage of CPE by western blotting and Coomassie blue staining after SDS-PAGE and identified a predominant CPE fragment only ∼1 kDa smaller than CPE and larger than the CPE fragment produced by trypsin. Since trypsin removes only the first 25 N-terminal amino acids from CPE, this result indicates that purified chymotrypsin does not affect CPE by removal of the 37 N-terminal amino acids, as previously reported [30].

Results using purified trypsin or chymotrypsin individually are informative but the intestinal lumen contains multiple proteases and some of those proteases have been shown to process other *C. perfringens* toxins. For example, serine proteases initiate processing of *C. perfringens* epsilon toxin, but carboxypeptidase then further processes that toxin [38]. Consequently, it was unclear whether, i] chymotrypsin or trypsin effects on CPE predominate in the intestines, ii] those two proteases act synergistically to process CPE beyond their individual cleavage effects, or iii] other intestinal proteases alone, or in combination with trypsin or chymotrypsin, cleave CPE beyond the individual effects observed for trypsin or chymotrypsin. Therefore, to better understand CPE processing during type F intestinal disease, the current study evaluated CPE processing by MIC *ex vivo* and in the small intestine.

The current results indicated that, when incubated *ex vivo* with MIC or added to the small intestine, CPE is mainly cleaved to a fragment matching the CPE fragment size generated by purified trypsin and ∼3 kDa less than native CPE. With some MIC preparations, and in the tested mouse small intestinal loops, a minor CPE band ∼7 kDa smaller than CPE was also observed, possibly suggesting there may be some animal-to-animal variability in CPE processing. It should also be noted that this minor ∼28 kDa CPE fragment is too small to be cytotoxic based upon results of previous CPE deletion analysis studies [42].

An important role for trypsin in CPE processing *ex vivo* received support from the ability of the trypsin inhibitor TLCK to affect this CPE processing by MIC. When MIC were preincubated with TLCK, there was an increase in the size and staining intensity of the main CPE fragment and staining intensity of the minor CPE also visibly decreased. Preincubation of MIC with chymotrypsin inhibitor did not affect normal CPE processing by MIC. When both trypsin and chymotrypsin inhibitors were present, CPE processing occurred similarly as in the presence of TLCK alone. Combining the pan-protease and trypsin-specific inhibitor results indicates that other intestinal proteases can affect CPE processing in the absence of trypsin although, as mentioned, trypsin is necessary and sufficient for processing CPE to the major fragment of ∼32 kDa. These inhibitor results also indicate that the minor processed CPE band of ∼28 kDa observed with some MIC preparations, and produced in the intestine, also involves trypsin activity. Of note, CPE structural biology studies [43, 44] showed that the N-terminal 34 amino acids of CPE [which includes the N-terminal trypsin cleavage site] have no detectable electron density by x-ray crystallography. Since this CPE region is disordered, it may be more accessible to intestinal proteases, including trypsin, when the toxin is free in solution.

Kinetically, the current study determined that CPE processing by MIC *ex vivo* is very rapid, reaching completion within 10 min. This rapid processing likely provides a major explanation for the detection of processed CPE in intestinal loops treated with CPE for only 15 min, i.e., considerable amounts of CPE may be proteolytically-processed even before this toxin binds to the intestines or becomes sequestered in large CPE complex. However, it remains possible that some CPE may be processed once it becomes localized in the large CPE complex given that CPE binding and large CPE complex formation also occur quickly *in vivo*, becoming detectable by the 15 min treatment time-point in mouse small intestinal loops. Consistent with the possibility that some CPE cleavage might occur after the toxin becomes sequestered in large complex, a previous study [45] showed that a portion of CPE in large complex remains exposed on the membrane surface, i.e., CPE antibodies react with CPE present in large complex in intestinal brush border membranes. If some processing of CPE localized in large complex does occur, this might involve the unstructured N-terminus of CPE remaining outside the large complex and therefore accessible to trypsin. It is also possible that CPE remains more susceptible to proteases when transiently located in small complex or before the prepore to pore shift. Given the absence of structural information concerning CPE complexes, these issues will await future studies.

Intestinal proteases vary in their effects on the toxicity of *C. perfringens* toxins produced in the intestines [1, 37, 38]. For some toxins [e.g., epsilon toxin] this processing has a toxicity-activating effect, but for other toxins [e.g., beta toxin] proteolytic cleavage is detrimental, with a toxin-inactivating effect. The current study first explored this issue *in vitro*. When CPE and MIC were preincubated together and that mix was then added to Caco-2 cells, this experiment revealed that CPE preincubated with MIC exhibits similar Caco-2 cell cytotoxicity as does CPE preincubated without MIC, i,e., proteolytic processing by MIC did not substantially affect CPE cytotoxicity. Consistent with the MIC-processed CPE retaining cytotoxic activity *in vitro*, the current study also demonstrated that, while much of the CPE associated with large complex formed in the intestines is proteolytically-processed, it retains intestinal enterotoxicity, i.e., in the presence of intestinal proteases, CPE still causes the intestinal damage important for type F disease [9, 28, 46].

Beyond providing important insights into CPE proteolytic processing under intestinal conditions, the current study also offers new information regarding large CPE complex formation and stability *in vivo*. The availability of mouse small intestinal loops treated for different times with CPE allowed this study to analyze large CPE complex formation kinetics and large CPE complex stability *in vivo*. CPE western blot studies showed that large CPE complex formation and, by extension, CPE binding [since it is required for large CPE complex formation [36]], occur rapidly in the intestines. These western blot results further indicate that by 30 min the vast majority of CPE injected into mouse small intestinal loops had already bound and become sequestered in large CPE complex. However, a significant percentage of the CPE present in large CPE complex after this 30 min treatment was still susceptible to dissociation by sample boiling prior to SDS-PAGE. In contrast, after 4 h of CPE treatment very little CPE could be released from the large complex by sample boiling. Collectively, these results indicate that the large CPE complex formed *in vivo* becomes more physically stable with time. Further studies are needed to better analyze how the formation and composition of large CPE complex influence stability of this complex *in vivo*.

## Materials and Methods

### Proteases and inhibitors

Purified trypsin [Bovine Pancreas TPCK-treated Trypsin] was purchased from Thermo Scientific, while purified chymotrypsin [Alpha Chymotrypsin TLCK-treated] was obtained from Worthington Biochemical Corporation. N-TOSYL-L-PHENYLALANINE CHLOROMETHYL KETONE [Synonym: TPCK] chymotrypsin inhibitor was purchased from Sigma, as was N-α-TOSYL-L-LYSINE CHLOROMETHYL KETONE [Synonym: TLCK] trypsin inhibitor. Protease Inhibitor Cocktail III Mammalian [AEBSF HCl: 100 mM, Aprotinin: 80 uM, Bestatin: 5 mM, E-64: 1.5 mM, Leupeptin: 2 mM, Pepstatin A: 1 mM] was purchased from RPI [Research Products International].

Clostridium perfringens *enterotoxin [CPE]*: As described previously [47] CPE was purified to homogeneity from type F strain NCTC8238 [ATCC 12916].

### Treatment of CPE with purified trypsin *in vitro*

Purified CPE [30 ng] was treated with 5, 10, 20, 30, 40, or 50 μg of trypsin in PBS [total volume 25 μl] for 10 min at 37°C. After this incubation, 5 μl of 5X loading buffer (which contains both SDS and β-mercaptoethanol) was added to each sample and the samples were then loaded onto a 12% acrylamide gel containing SDS. After electrophoresis, the separated proteins on these gels were electrotransferred onto nitrocellulose membranes [0.45 μm pore size, Biorad], which were then blocked with 5% milk in Tris-buffer saline with 0.1% Tween-20 [TBST]. The blocked membranes were incubated overnight at 4°C with a 1:1000 dilution of polyclonal CPE antibody [48]. On the next day, the blot was incubated with a 1:10000 dilution of goat anti-rabbit IgG antibody conjugated with horseradish peroxidase [Invitrogen] at RT for 1 hour with gentle shaking. The blot was developed using SuperSignal West Pico PLUS Chemiluminescent Substrate [Thermo Scientific].

The above experiment was also performed using a reaction containing 5 μg of CPE in PBS [total volume 25 μl] with 5-50 μg of trypsin in PBS that was incubated for 10 min at 37°C, followed by electrophoresis, as above, and staining the gels with Coomassie blue [G-250, Biorad].

### Treatment of CPE with purified chymotrypsin *in vitro*

Purified CPE [30 ng] was treated with 5, 10, 20, 30, 40, or 50 μg of chymotrypsin in PBS [total volume 25 μl] for 10 min at 37°C. After this treatment, CPE western blotting was performed as described above for trypsin treatment. Similarly, the above experiment was repeated using a reaction containing 5 μg CPE in 25 μl of PBS with 5-50 μg of purified chymotrypsin that was incubated for 10 min at 37°C, followed by electrophoresis [as described above] and staining the gels with Coomassie blue stain.

### Preparation of MIC

Small intestinal lumen contents were obtained from healthy ∼4 month old BALB/c mice after they had been euthanized by CO_2_ asphyxiation for colony management [IACUC protocol 23482]. Equal numbers of male and female mice were used. After euthanasia, small intestinal lumen contents were immediately harvested from excised small intestine by gentle squeezing and stored at −80°C. For use in experiments, each intestinal content was thawed, weighed and mixed with PBS buffer at a ratio of 1:1. The mixture was then centrifuged for 5 min and supernatants [now designated as MIC] were collected, aliquoted, and stored at −80°C until use in the current study. Typically, one MIC preparation was used for each experimental repetition.

### Comparison of CPE processing by MIC from different mice

To evaluate *ex vivo* effects of MIC from different mice on CPE, 30 ng of native CPE were incubated in 25 μl of PBS with 1 μl of MIC from three different mice [2 female and 1 male] for 10 min at 37 °C. After incubation, 5 μl of 5X loading buffer was added to each sample and the samples were then boiled for 5 min before loading onto a 12% acrylamide gel containing SDS. After electrophoresis, separated proteins were electrotransferred onto a 0.45 μm pore size nitrocellulose membrane and processed for CPE western blot analysis as described earlier.

### Comparison of CPE processing by MIC, purified trypsin or purified chymotrypsin

For CPE western blot analyses, 30 ng of native CPE was added to PBS containing 1 μl of MIC, 20 μg of purified trypsin or 20 μg of purified chymotrypsin [reaction volumes 25 μl] for 10 min at 37°C. Identical MIC, trypsin or chymotrypsin samples were similarly incubated without CPE. After this incubation, all samples were treated with 5 μl of 5X loading buffer and this sample was then boiled for 5 min; the trypsin- or chymotrypsin-treated samples were not boiled to reduce CPE aggregation. All samples were then electrophoresed on 12% acrylamide gels containing SDS and processed for CPE western blotting, as described earlier. For Coomassie blue staining analyses, 5 μg of native CPE was incubated in PBS [reaction volume of 25 μl] containing 2.5 μl of MIC, 20 μg of purified trypsin or 20 μg of purified chymotrypsin. The samples were then processed and electrophoresed on a 12% acrylamide gel containing SDS as described above for CPE western blots. After electrophoresis, gels were stained with Coomassie blue.

### Effects of protease inhibitors on MIC processing of CPE ex vivo

Purified trypsin [10 μg] was preincubated with 200 μM TLCK or 200 μM TPCK in PBS [total reaction volume was 20 μl], while 10 μg of purified chymotrypsin was preincubated with 200 μM TPCK or 200 μM TLCK in PBS [total reaction volume was 20 μl]. A control sample contained only 20 μl of PBS. All samples were preincubated at 37°C for 30 min. After this preincubation, 5 μg of CPE in 5 μl of PBS was added to each sample before incubation for 10 min at 37°C. The incubated samples were then treated with 5 μl of 5X loading buffer and electrophoresed on 12% acrylamide gels containing SDS, followed by staining those gels with Coomassie blue. For MIC treatments, MIC [1 μl] in PBS [total reaction volume of 20 μl] was preincubated with 200 μM TLCK, 200 μM TPCK, 200 μM each of TCPK and TLCK, or 5 μl of Protease Inhibitor Cocktail III. As a control, 1 μl of MIC was preincubated in 20 μl of PBS. These preincubations were performed at 37°C for 30 min. After this preincubation, 5 μl of PBS containing 5 μg of CPE was added to each sample and the samples were incubated for 10 min at 37°C. These samples were then treated with 5 μl of 5X loading buffer, boiled for 5 min, and then electrophoresed on 12% acrylamide gels containing SDS, followed by staining the gels with Coomassie blue staining.

### Caco-2 cell culture

Caco-2 cells [ATCC] were grown in Eagles Essential Medium [Lonza] supplemented with 10% fetal bovine serum [Gibco], 1% MEM nonessential amino acids [Cytiva], 100 μg/ml of penicillin-streptomycin solution [Corning] and 1% glutamine [Corning]. These cultures were maintained at 37°C in a 5% CO_2_ atmosphere.

### MIC effects on large CPE complex formation by Caco-2 cells

Caco-2 cells were grown in 6 well cell culture dishes for 5-6 days. Cultures were then treated for 15 min at 37°C with CPE [0.5 μg/ml] in the presence of 1 μl of MIC in 1 ml of HBSS; the MIC was first added into the culture, immediately followed by the addition of CPE. Other Caco-2 cell wells were treated with 1 ml of HBSS alone or with 1 ml of HBSS containing 1 μl of MIC [no CPE]. After a 15 or 30 min incubation, each sample was gently collected using a cell scrapper and lysed with RIPA buffer [Boston BioProducts, Inc.] in the presence of Benzonase Nuclease [Millipore] and Protease Inhibitor Cocktail III at RT for 30 min with gentle absorption. The samples were centrifuged and the lysates were collected in sterile tubes. A 1 μl aliquot of lysate from each treatment was then mixed with 24 μl of HBSS buffer plus 5 μl of 5X loading buffer and the total 30 μl final volume was loaded onto a 6% SDS PAGE gel and processed for CPE western blotting as already described.

### Effects of MIC pretreatment on CPE cytotoxicity and large CPE complex formation for Caco-2 cells

HBSS containing native CPE [3 μg] in the presence or absence of 2.5 μl of MIC, HBSS containing 2.5 μl of MIC, or HBSS alone [total sample volumes of 25 μl] were preincubated at 37°C for 10 min. After that preincubation, samples were adjusted with HBSS to a 1 μg/ml CPE concentration, with a similar dilution of the MIC alone [no CPE] sample. Each sample [1 ml] was then added onto confluent Caco-2 cell cultures for 1 h at 37°C. After this treatment, culture supernatants were collected, centrifuged and the lysates were processed for LDH-release, as an indicator of cytotoxicity, using the CyQUANT LDH Cytotoxicity Assay Kit [Invitrogen].

Adherent cells in these cultures were gently collected using a cell scrapper and lysed with RIPA buffer with Benzonase Nuclease and Protease Inhibitor Cocktail III at RT for 30 min. The samples were centrifuged and a 1 μl aliquot of lysate from each treatment was then mixed with 24 μl of HBSS buffer plus 5 μl of 5X loading buffer; the total final volume was then loaded onto a 6% acrylamide gel with SDS and processed for CPE western blotting as already described.

### CPE treatment of mouse small intestinal loops

To assess the effects of intestinal proteases on *in vivo* CPE processing, large CPE complex formation and enterotoxicity, the mouse small intestinal loop model was used. Balb/C mouse small intestine segments [∼10 cm] were surgically ligated under general anesthesia, as previously described [26]. These intestinal loop experiments were conducted using 6 groups of mice [*n* = 6 per group, with equal numbers of male and female mice]. Each loop was challenged with 1 ml of HBSS that did or did not contain 100 μg of purified CPE. After 15 min, 30 min or 4 h of incubation, the mice were euthanized. Samples from these treated intestinal loops were collected, fixed in buffered formalin [10%], pH 7.2 for 72h and then processed to 4 μm sections before staining with heamatoxylin and eosin [HCE]. These samples were examined by a pathologist in a blinded fashion. Contents of the intestinal loops were collected by gentle squeezing and stored at −80°C until use in experiments. These *in vivo* experiments were reviewed and approved by the University of California Davis Institutional Animal Care and Use Committee [Protocol 23182].

### Analysis of large CPE complex formation and stability in vivo

Intestinal contents from control or CPE treated mice, as described above, were centrifuged and half of the supernatant volume was boiled for 5 min while the other half was not. These samples were then electrophoresed on 12% acrylamide gels with SDS, or on 6% acrylamide gel with SDS, and processed for CPE western blotting as described above for Caco-2 cells. Samples run on 12% acrylamide gels with SDS were boiled for 5 min before electrophoresis and western blotting.

Statistical analyses. The statistical significance of CPE cytotoxicity results was assessed by one way ANOVA with Dunnett’s multiple comparison test.

## Acknowledgements

This study was generously supported by grant R01AI019844-41 from the National Institute of Allergy and Infectious Diseases. The content is solely the responsibility of the authors and does not necessarily represent the official views of the National Institutes of Health.

## Institutional Review Board Statement

Not applicable.

## Informed Consent Statement

Not applicable.

## Data Availability Statement

The authors confirm that the data supporting the findings of this study are available within the article.

